# The Evolutionary Dynamics of Cooperation in Collective Search

**DOI:** 10.1101/538447

**Authors:** Alan N. Tump, Charley M. Wu, Imen Bouhlel, Robert L. Goldstone

## Abstract

How does cooperation arise in an evolutionary context? We approach this problem using a collective search paradigm where interactions are dynamic and there is competition for rewards. Using evolutionary simulations, we find that the unconditional sharing of information can be an evolutionary advantageous strategy without the need for conditional strategies or explicit reciprocation. Shared information acts as a recruitment signal and facilitates the formation of a self-organized group. Thus, the improved search efficiency of the collective bestows byproduct benefits onto the original sharer. A key mechanism is a visibility radius, where individuals have unconditional access to information about neighbors within a limited distance. Our results show that for a variety of initial conditions—including populations initially devoid of prosocial individuals—and across both static and dynamic fitness landscapes, we find strong selection pressure to evolve unconditional sharing.

## Introduction

Social behavior is structured by the dynamics of the environment and how we interact with one another. Strategies that thrive in one context may be poorly suited to others. How do social behaviors arise in an evolutionary context? And can the dynamics of social interactions support the emergence of cooperation without appealing to conditional strategies?

Evolution is often summarized as “survival of the fittest”, evoking a notion of fierce competition between individuals. Where is there room for prosociality and cooperation in the midst of evolutionary competition? One of the early challenges for Darwin’s theory of evolution (1859) was to explain the origin of prosocial adaptations that improve the welfare of others or one’s group as a whole, but at a potential cost to the individual. Darwin’s explanation appealed to the notion of *group selection*, where the costs of altruism are ultimately justified by increased fitness for the group (Darwin, 1871). Thus, groups with more prosocial members may outcompete rival groups. Although group selection offers a potential pathway for the emergence of cooperation, it often requires strong assumptions, such as stable group structures and strong competition between groups (Janssen & Goldstone, 2006). Without these assumptions, selection at the individual level can undermine group selection. Thus, a comprehensive understanding of prosociality requires a theory of individual selection (Wilson & Wilson, 2007).

### Theories of Cooperation

One traditional explanation for individual selection of prosociality is through the mechanism of *kin selection* (also known as inclusive fitness), where recipients of altruistic acts tend to be genetically related to the donor (Nowak, 2006). Hamilton’s law (1964) states that the costs of prosociality *C* must be justified relative to the benefits of the recipient *B* by accounting for the relatedness of individuals *r* such that 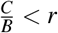. While kin selection explains prosociality between genetically similar individuals, Hamilton’s law alone fails to account for all the social behaviors we see in human society (Rand & Nowak, 2013; Fehr & Fischbacher, 2003) and in animals (e.g., Spottiswoode, Begg, & Begg, 2016; Brown, Brown, & Shaffer, 1991). Many mechanisms have been proposed in order to justify the evolution of cooperation towards nonrelatives, typically requiring an initial investment of a donor towards a non-related individual with expectations of reciprocity or benefits.

#### Conditional Cooperation

Theories of conditional cooperation operate on expectations of future reciprocity, where seemingly prosocial behavior is ultimately grounded in self-interest. Often described as impure altruism (Andreoni, 1989), both direct and indirect reciprocity appeal to conditional strategies (e.g., tit for tat; Nowak & Sigmund, 1992), where individuals conditionally cooperate with each other, so long as future reciprocation is expected. Direct reciprocity depends on multiple interactions with the same individual, while indirect reciprocity typically relies on reputation systems, where cooperative behavior is used as a social signal to third-parties (Nowak & Roch, 2007). Conditional cooperation has been widely studied in the context of game theory, yet simple mechanisms of social or spatial dynamics can also explain the origins of cooperation (Nowak & May, 1992).

#### Unconditional Cooperation

Theories of unconditional cooperation explain the origin of prosocial behavior through changes in the interaction structure for the donor (Perc, Gómez-Gardeñes, Szolnoki, Floría, & Moreno, 2013). Thus, behaving prosocially can make it more likely to interact with other prosocial individuals. *Network reciprocity* operates on similar principles as kin selection, but where the cost-benefit ratio is defined relative to interaction partners (Nowak, 2006). This approach has shown that by situating agents on a network (Ohtsuki, Hauert, Lieberman, & Nowak, 2006) or in a spatial landscape (Nowak & May, 1992), prosocial individuals tend to interact more with similar partners, thus creating self-organized regions where prosociality proliferates (Perc et al., 2013). It is also possible to replace spatial similarity or network connectivity with some arbitrary feature or tag (Riolo, Cohen, & Axelrod, 2001), such that individuals with similar features are more likely to interact with one another. This provides a useful bridge between individual and group level mechanisms, because it describes how groups can form based on spatial, network, or feature similarity.

Two key assumption are made by these theories. The first is that the initial population already includes multiple prosocial individuals (Nowak & May, 1992; Ohtsuki et al., 2006). Yet this doesn’t answer the crucial question of how cooperation emerges *ex nihilo*. Secondly, the interaction structures are more or less static: agents are either embedded in some spatial location (Nowak & May, 1992), as a fixed node in a network (Ohtsuki et al., 2006; Barkoczi, Analytis, & Wu, 2016), or given a fixed feature tag (Riolo et al., 2001). While groups can still emerge through the dynamics of evolution, interaction partners remain relatively stationary (but see Janssen & Goldstone, 2006) and individual dynamics (e.g., search behavior) are largely unaccounted for.

*Pseudo-reciprocity* is a related theory of unconditional cooperation, where the key difference from network reciprocity is that the fitness of the donor does not depend on the phenotype of the recipient. Thus, prosocial behavior can be beneficial without depending on the presence of other prosocial individuals in a group. Prosociality can alter the social environment for the donor (e.g., by sharing information about resources), such that the donor gains byproduct benefits through self-interested behavior of the recipients (Connor, 1986; Brown et al., 1991). For example, Cliff Swallows (*Hirundo pyrrhonota*) share information about the location of insect swarms through a unique vocal signal (i.e., a food call), which attracts other peers. While it is difficult to track the insect swarms individually, the collective recruited by the information sharer tracks the swarm more efficiently. Hence, even without expectations of reciprocity (i.e., future vocal signals from peers), each individual benefits by behaving prosocially and sharing information (Brown et al., 1991). Thus, pseudoreciprocity offers a mechanism where individuals can be unconditionally prosocial towards all the members of the group, rather than towards a restricted set of cooperative partners.

### Goals and Scope

Here, we analyze the emergence of cooperation through sharing information. We use evolutionary simulations to study how individual selection pressure can give rise to sharing, even from initial populations void of prosocial individuals. We simulate agents searching for rewards on a high dimensional fitness landscape, where the flow of information is dynamically and spatially defined. Agents have a binary phenotype that defines whether or not they share information unconditionally to the rest of the population. We show that this global sharing signal acts as a recruitment mechanism that facilitates the self-organization of dynamic groups. Because groups are more effective at finding rewards than lone individuals, we find that sharing emerges and dominates our evolved populations across a large range of initial conditions and in both static and dynamic fitness landscapes.

## Collective Search Simulations

We use a multi-agent framework based on Bouhlel, Wu, Hanaki, and Goldstone (2018), who found that sharing information can be beneficial to the donor, even in competitive contexts and without expectations of reciprocity. The costs of sharing information (through resources lost to competition) can be outweighed by the byproduct benefits of cooperation. A simple coordination mechanism of a local visibility radius (i.e., nearby agents have access to each others’ rewards) facilitates the formation of a self-organized collective. Thus, sharing information acts as a recruitment signal, attracting others to the donor, and increasing the likelihood of future social interactions (via the visibility radius). These future interactions are the source of byproducts benefits for the sharer. Here, we use evolutionary simulations and more extreme levels of competition (compared to Bouhlel et al., 2018) in order to study how sharing interacts with innovation, and under which initial conditions there exists individual selection pressure for unconditional sharing, leading to group-level cooperation (Goldstone & Janssen, 2005). Code for reproducing these results is publicly available at https://github.com/alantump/adaptiveSharingEvolution.

### Methods

Adopted from Bouhlel et al. (2018), we simulate groups of *k* agents searching for rewards on a 10-dimensional^1^ fitness landscape over *T* = 50 trials. On each trial *t*, agents can use either individual or social information (see below) to search for rewards on the fitness landscape. Payoffs are proportional to the inverse Manhattan distance of agent *i* from a global optimum Ω:

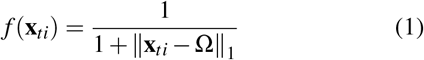

where **x***_ti_* contains the coordinates for each dimension *m* = 1,…, 10 of the current location of agent *i* at trial *t*. The coordinates of the global optimum Ω are sampled from a uniform distribution 𝒰(1,10) for each dimension.

#### Competition

The payoffs *f*(**x***_ti_*) are subject to competition, which we implement by having agents split rewards when occupying nearby spaces in the environment. Specifically, we use a competition parameter *c* that defines an exponentially decaying competition metric *C*(**x***_ti_*, **x***_t j_*) between each pair of agents *i* and *j*:

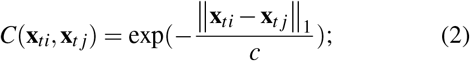

Larger values of *c* induce higher competition over larger distances (see Fig. 1a), while in the limit of *c* → 0, competition only occurs when agents occupy the exact same solution (as in Bouhlel et al., 2018). Splitting of rewards is proportional to the sum of competition values for all other agents. Hence, for location **x***_ti_*, the acquired reward is:

**Figure 1:**
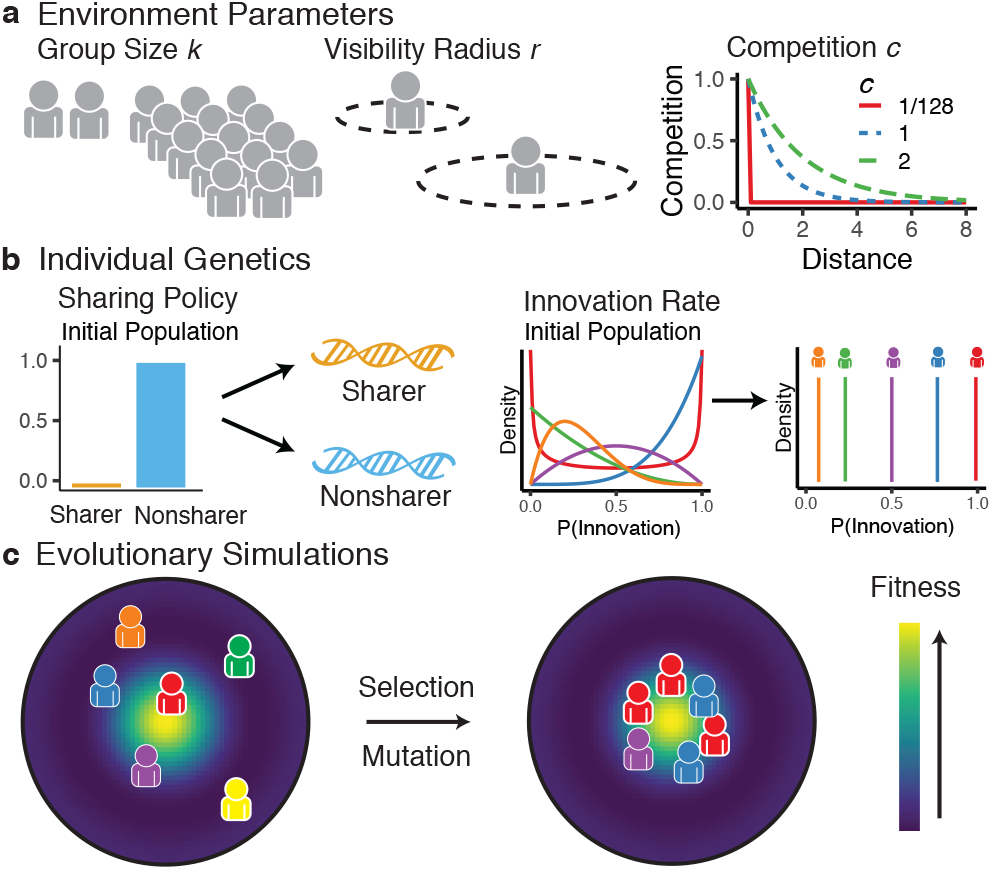
Evolutionary Simulations. **a**) We vary three main environmental parameters: group size, the visibility radius, and competition level. Group size *k* specifies the number of agents interacting together. Visibility radius *r* defines the maximum Chebyshev distance between two agents where information can be passively observed. Competition level *c* defines the decay rate of an exponential competition function that determines how agents split rewards (higher values of *c* result in splitting over larger distances. **b**) Each agent is defined by a sharing policy (either sharer or non-sharer) and an innovation rate (between 0 and 1). **c**) We use evolutionary simulations over 200 generations to see which individual genes emerge through selection pressure and mutation.

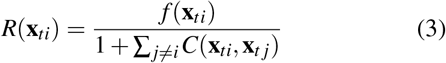

#### Individual search

Each agent begins at a random starting location, where each dimension is sampled from a uniform distribution 𝒰(1,10). On every trial, each agent *i* stores the location **x***_tj_* and reward value R(**x***_ti_*) of both individually and socially acquired information (see information sharing and visibility radius). We use a local search strategy, where the agent selects the location with the largest observed reward value 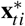 up until time *t*, and then has an opportunity to innovate on it by modifying each value in 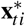 by a discrete value in {−1,0,1}.

We define the *Innovation rate* as the probability that an agent innovates, where otherwise 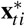 is copied verbatim. If the agent innovates, we modify each dimension of 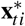 by drawing from a Binomial distribution centered on zero 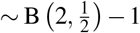. Intuitively, half of the time there is no change along that dimension, while changes of both −1 or +1 are equally likely, each with a probability of 25%.

#### Social information

Depending on their sharing policy, agents are deterministically either sharers or non-sharers. Sharers will unconditionally share information about both reward location **x***_ti_* and value R(**x***_ti_*) to all other agents, while non-sharers will withhold it. Sharing information is associated with an increased cost due to splitting rewards with imitators, but can also confer byproduct benefits by broadcasting high quality solutions, which are subsequently modified by group members and improved upon, before being transmitted back via the visibility radius or by other sharers.

In addition to the global sharing signal, we use a visibility radius as a feature of the environment. At each trial *t*, agents passively provide information about reward locations and magnitudes to other agents that are within visibility radius *r*. For any two agents *i* ≠ *j*, agent *j* is visible to agent *i* if the maximal distance between the two agents on any dimension (i.e., the Chebyshev distance) is not greater than the visibility radius *r*:

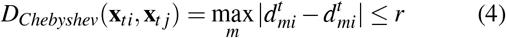

The visibility radius is a coordination mechanism that allows for localized transmission of information. Whereas the sharing signal is a global mechanism operating at all distances, the local visibility radius allows for dynamic interaction structures to emerge and facilitates the spontaneous formation of spatially coherent groups. Crucially, given the high dimensionality and size of the search space, it is unlikely for any two agents to fall within the same visibility radius without explicit information sharing. For example, there is 0.1% probability of two agent being visible to one another at initialization for a radius of 2.

### Evolutionary Simulations

Inspired by biological evolution, we embed the simulation framework in an evolutionary algorithm, which uses selection pressure and mutations over multiple generations to discover which sets of behavioral parameters evolve. The evolutionary algorithm is well suited for our research question because fitness-maximizing behavior (e.g., willingness to share information) depends on the behavior of others in a game theoretical context.

#### Initial conditions

Beginning with a population of 300 agents, each agent carries genes determining innovation rate and sharing policy (i.e., sharer or non-sharer). We start with an initial population consisting of only non-sharers to address the question of how cooperative behavior can emerge *ex nihilo* through individual selection. We vary the initial mean innovation rate in the populations to ensure that the results of the evolutionary algorithm are not dependent on the starting conditions. The initial values for innovation rate were sampled from a Beta distribution, with the mean of the distribution sampled from a uniform distribution 𝒰(0, 1).

For each generation, we repeatedly sample *k* agents from the whole population. We simulate these agents performing collective search, where behavior is determined by their genetic makeup (innovation rate and sharing policy). We repeat the simulation procedure over 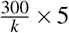 repetitions, resulting in approximately 5 simulations per agent in each generation.

#### Selection and mutation

We select the agents with the highest fitness to produce genetically similar offspring via tournament selection. In this selection procedure, we repeatedly sample 7 random individuals from the population, whereby the individual with the highest relative performance passes its genes onto the next generation. This selection process is repeated 300 times in order to produce a new generation of 300 agents. The genes of the new generation are exposed to weak mutation to consistently ensure gene variation, where each gene has a probability of mutation. The sharing gene mutates with *p* = .002, whereby a new sharing policy is drawn from a binomial distribution 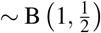, with the new policy equally likely to be sharer or non-sharer. The innovation gene mutates with *p* = .02, whereby the previous innovation is modified by adding Gaussian noise ~ N (0,0.2). The innovation rate was truncated between [0,1]. Note that we chose the mutation probabilities and strengths to be high enough to ensured constant variation in the gen pool.

The genetic algorithm repeats the process of fitness evaluation, selection, and then reproduction with mutation for 200 generations to ensure the population converges to a stable outcome. We ran 10 replications of this procedure and report the average evolved parameters of the last 10 generations (i.e., generations 190 to 200) over each of the 10 replications. We systematically varied group size (*k* ∈ [2,… 14]), visibilityradius (*r* ∈ [0,1,2,3,4]), and competition level (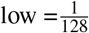; medium = 1; high = 2) to investigate how the structure of the environment influences the selection of individual characteristics (sharing and innovation).

## Results

When exposed to selection pressure via the evolutionary algorithm, the populations evolved different sharing and innovation rates depending on the environmental parameters (see Fig. 2 for examples). Figure 3 shows the proportion of sharers and the innovation rate at equilibrium for different parameter combinations, where yellow tiles indicate high levels of either sharing or innovation.

**Figure 2:**
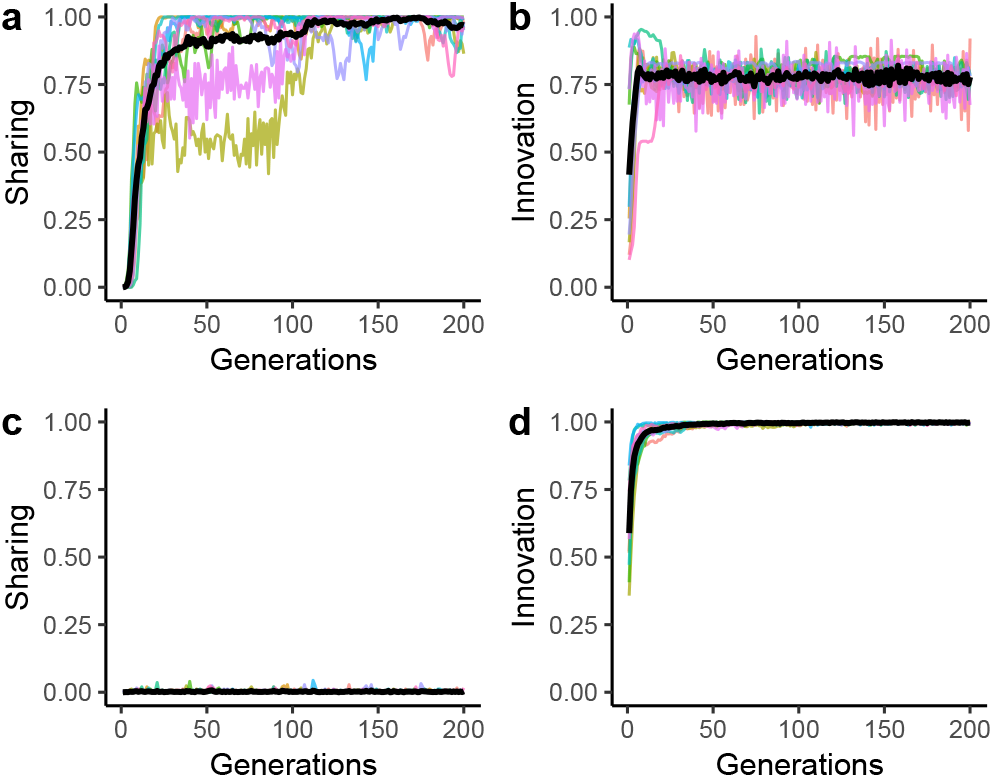
Evolution of sharing and innovation over 200 generations. **a-b**) An example where populations evolve high sharing and innovation rates, with group size *k =* 6, visibility radius *r =* 4 and competition level *c* = 1/128. **c-d**) An example where individuals adopted high innovation rates but did not evolve sharing, based on group size *k* = 6, visibility radius *r* = 0 and competition level *c* = 2. Each colored line represents the average parameter value within a population, while the black line indicates the average across populations.

**Figure 3:**
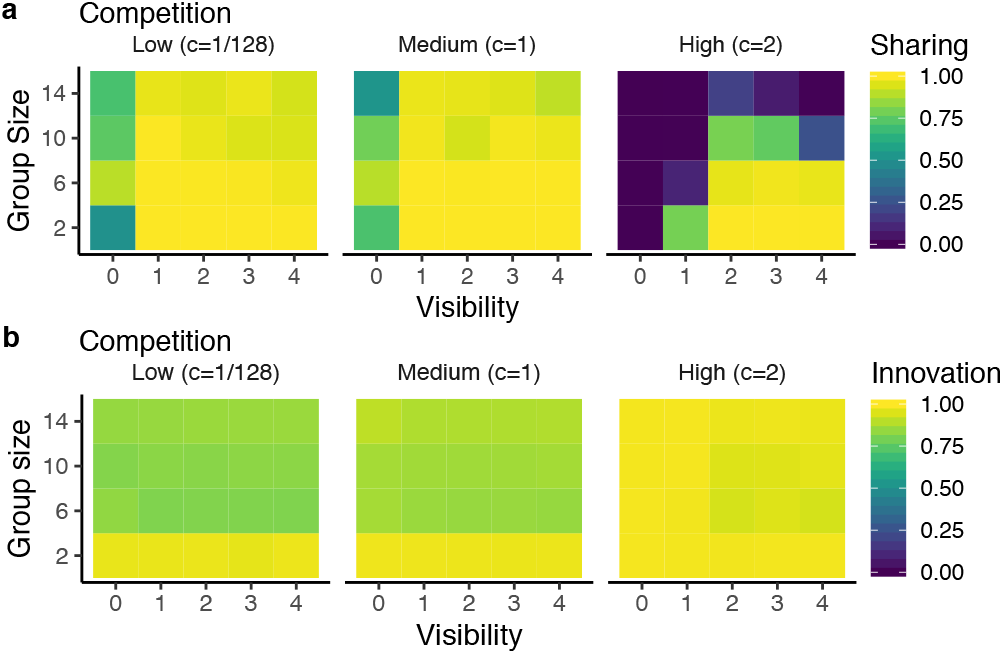
Equilibrium results for different combinations of environmental parameters. **a**) Agents evolved high sharing rates in low and medium competitive environments, although sharing was found in more restricted contexts under high competition (requiring smaller groups and larger visibility radius values). **b**) Overall, we find high levels of innovation, although we also see the trend that larger groups evolve slightly lower innovation rates.

### Sharing evolves ex nihilo

Starting from initial conditions of no sharers in the population, we find that sharing emerges in the overwhelming majority of our simulation parameters, and that sharing often dominates the population at close to ceiling levels (Fig. 3a). However, we also discover the limits of sharing as an adaptive strategy as we increase the level of competition for rewards. Under high levels of competition, only smaller groups with larger visibility radius are able to support sharing.

### Sharing and innovation co-evolve

We find that over the entire parameter space, all populations evolved high innovation rates (Fig. 3b), although not at ceiling level (i.e., yellow tiles) compared to sharing behavior. Looking more closely, we find relatively higher innovation rates in small groups compared to large groups, with this effect most pronounced under low or medium levels of competition. Yet, how are sharing and innovation behaviors related to each other?

To further understand the interaction between strategies, we ran additional simulations with innovation rate fixed at low (25%), medium (50%) or high (100%) values. The results are shown in Figure 4, where we replicate the main findings of the previous simulation for high innovation rate (top row). However, we find that sharing becomes substantially less adaptive for populations with innovation fixed at low or medium levels. Thus, innovation is an essential ingredient for prosocial traits to develop, as has been shown in previous work on cultural transmission through iterative cycles of imitation and innovation (Ehn & Laland, 2012; Wisdom & Goldstone, 2011; Derex, Feron, Godelle, & Raymond, 2015).

**Figure 4:**
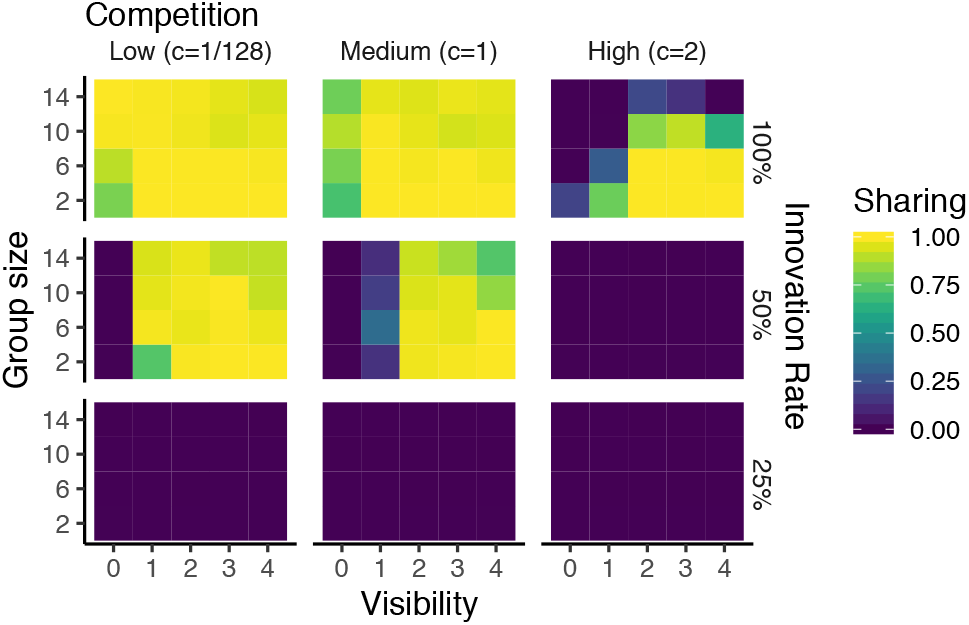
Equilibrium results of sharing for fixed innovation rates (rows) at various parameter combinations. When the innovation rate is fixed at 100%, we largely replicate the results in Figure 3. However, when innovation is fixed at 50%, we find that sharing evolves in a more restricted set of parameters and exclusively with a visibility radius of 1 or larger. When there is an innovation rate of 25%, we find virtually no emergence of sharing.

### Interim conclusion

We show that sharing can evolve across a variety of different environments and in mixed groups with different proportions of sharers and non-sharers. The selection pressure for sharing can lead to it becoming a dominant trait prevalent in the vast majority of the population. The spatial dynamics of this simulation framework facilitated by a visibility radius lead to a setting where selection pressure does not prioritize freeriding and the group does not succumb to a tragedy of the commons.

## Dynamic Simulations

We now extend the framework to account for a changing environment, implemented by a wandering global optima. We define the global optima Ω^*t*^ and modify it on each time *t* with a probability determined by the environmental change rate *p_e_*. With probability *p_e_*, the environment’s global optima changes, otherwise it stays the same (Ω^*t*+1^ = Ω^*t*^). When the environment changes, each coordinate of the global optima 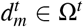 has a 50% probability of being modified by +1 or −1, and a 50% probability of remaining the same. This is the same as the local search rule used by individual agents.

In order to account for the decreasing validity of past observations in a changing environment, we introduce a temporal discount rate γ. Thus, the history of past observations maintained by each agent decays as a function of the elapsed time:

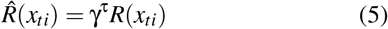

where 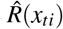 is the discounted reward and τ is the elapsed time between the observation and the current time. Thus, agents locally search around the reward location that has the largest discounted reward 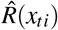. Both individually and socially acquired information follow the same decay rate. In our simulations we fixed the Discount rate γ = .99, which approximately corresponds to a 10% discount after 10 trials.

### Dynamic Results

Figure 5 shows the equilibrium results of our dynamic simulations. Again, we find that sharing is a beneficial strategy under many environmental conditions (Fig. 5a). Similar to the static case, there are limits to the conditions under which sharing emerges, particularly in highly competitive environments. The relationship between the visibility radius and group size becomes increasingly important, where a larger radius allows sharing to emerge in larger groups. We also observe that the evolved proportion of sharers decreases in more volatile environments (higher change rates) and in larger groups. This interaction is not observed in the static environment, but may be partially due to the increased difficulty of coordination and because out-of-date information can harm instead of help others (Boyd & Richerson, 1988; Henrich & Boyd, 1998). Additionally, we find that environmental change increases the evolved innovation rates (Fig. 5b). The intermediate levels of innovation found in the static simulations are eclipsed by even higher rates under environmental change.

**Figure 5:**
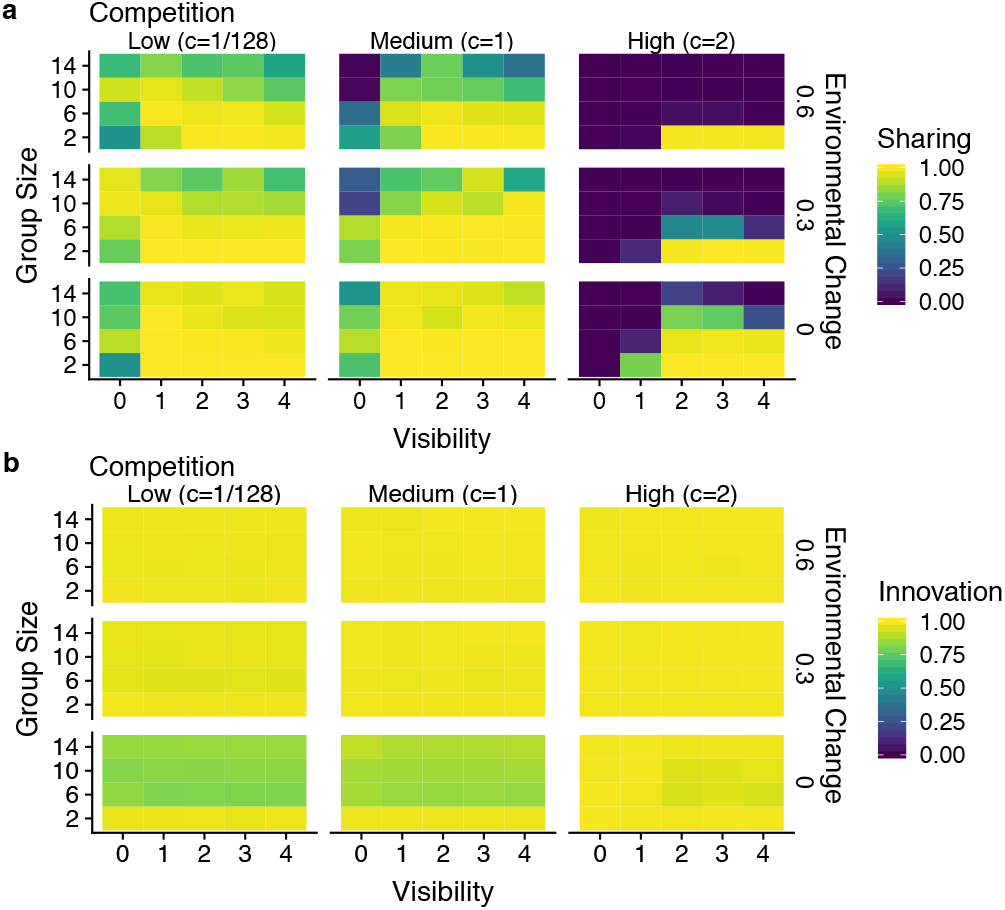
Equilibrium results in a dynamic environment. **a**) Again we find high sharing rates in low and medium competitive environments, but now higher rates of environmental change reduced the levels of sharing in the population. We also see stronger indications of an interaction between group size and visibility radius, where a larger visibility radius is required to coordinate larger groups and support a larger sharing population. **b**) Across all parameters, we find high levels of innovation emerge, although lower competition and larger groups reduces the extent of innovation.

## General Discussion

We use evolutionary simulations to show that for a variety of initial conditions and across both static and dynamic fitness landscapes, there exists individual selection pressure for the unconditional sharing of information. To summarize the effects of each environmental parameter on the equilibrium characteristics of innovation and sharing, we fit a linear model on the dynamic simulation results (Fig. 6). The size of the visibility radius contributes positively to the rate of sharers in the evolved population, while group size, environmental change, and competition all reduce the rate of sharers. Thus, the evolution of cooperation in the absence of reciprocity operates at a fine balance between coordination (via the visibility radius) and discord (through competition and the communication of out-of-date information).

**Figure 6:**
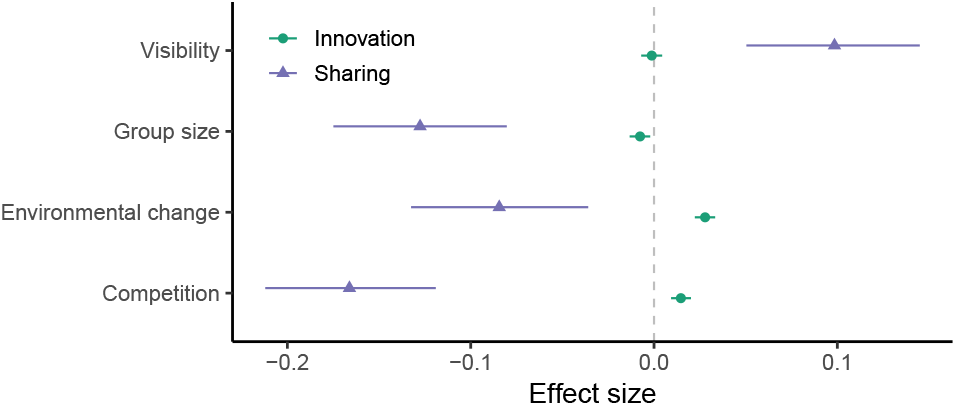
Regression results. The estimated effect sizes of environmental parameters on innovation rate (green) and sharing (purple). Error bars show 95% CI.

In comparison, the environmental effects on innovation are relatively small. We find relatively high levels of innovation in all simulations. Environmental change had the strongest influence on innovation, while higher competition also increased innovation. Rather, the more interesting result of our simulations involves the interaction between innovation and sharing, which co-evolve and are dependent on one another for producing the emergent behavior of collective search.

### How do the dynamics of cooperation work?

To get a deeper understanding of how sharing improves the welfare of the donor, we present a vignette of an agent who is either a sharer or a non-sharer in a population of non-sharers (Fig. 7). The sharer transmits a global signal that recruits peers and gathers them within visible range (Fig. 7a, orange line). This means that a sharer will have access to more social information compared to a non-sharer by being closer to others (Fig. 7a, blue line). Since we find high rates of innovation in all simulations, any imitated information is also tweaked and modified. Some of these modifications will improve upon the originally copied solution. This creates a feedback cycle of solutions that are consistently improved over time, which can benefit the original sharer through local transmissions within the visibility radius (Fig. 7b). Compared to a group of non-sharers (blue line), the sharer is able to explore the reward landscape better and achieve higher rewards despite the stronger local competition (Fig. 7c, orange line).

**Figure 7:**
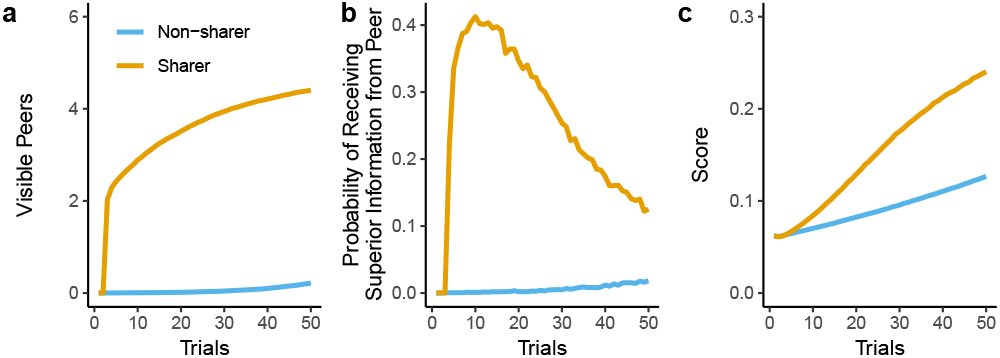
How sharing leads to cooperation. These results are the mean performance over 10,000 replications with a group size of *k* = 6, a visibility radius of *r* = 2, a innovation rate of 1 and a competition level *c* = 1. **a**) The sharer (orange line) attracts other individuals within their visibility radius through the sharing signal, leading to richer informational exchanges than compared to a non-sharer (blue line). **b**) Individuals who imitate the shared information also innovate, and thus passively provide improved information to the sharer through the visibility radius. **c**) As a result, the sharer benefits from passively gained information and acquires an overall higher pay-off compared to individuals in a non-sharing group.

In summary, as is the case with the Cliff Swallows (Brown et al., 1991), sharers recruit peers within their visibility radius and reap the byproduct benefits of passively acquired modifications to the original solution. Intuitively, larger visibility increases the ability of a group to stay connected with one another. However, the global sharing signal is an essential recruitment device that facilitates the formation of a group in the first place. Group coherency facilitated by the visibility radius provides byproduct benefits to the originator of the sharing signal, creating a feedback loop of imitation with innovation.

## Conclusion

Through the lens of evolution, we show how individual selection pressure can give rise to the unconditional sharing of information. The sharing signal does not require expectations of reciprocity in order to be beneficial, but rather directly benefits the sharer through the byproducts of cooperation. Shared information about a high reward acts as a recruitment signal, which leads to the emergence of a self-organized collective centered on the original donor. A key ingredient is a visibility radius, which allows individuals to observe the rewards of neighbors within a fixed spatial distance. This visibility radius provides a simple yet effective coordination mechanism that is grounded in simple spatial and social dynamics, creating complex patterns of emergent behavior.

More broadly, our results indicate that prosocial behaviour can evolve from initial conditions devoid of other prosocial individuals. While theories explaining the evolution of conditional reciprocity have been very influential (Nowak & May, 1992; Ohtsuki et al., 2006), our results provide an explanation for the initial emergence of prosocial individuals, which is an essential requirement for both conditional cooperation and group selection. Future implementation of conditional strategies in our framework could provide further insight into how various strategies co-evolve.

1 Bouhlel et al. (2018) studied environments of different dimensionality, while here we use 10-dimensional environments as a prototypical example.

